# Proteogenomics of adenosine-to-inosine RNA editing in the fruit fly

**DOI:** 10.1101/101949

**Authors:** Ksenia G. Kuznetsova, Anna A. Kliuchnikova, Irina I. Ilina, Alexey L. Chernobrovkin, Svetlana E. Novikova, Tatyana E. Farafonova, Dmitry S. Karpov, Mark V. Ivanov, Anton O. Goncharov, Ekaterina V. Ilgisonis, Olga E. Voronko, Shamsudin S. Nasaev, Victor G. Zgoda, Roman A. Zubarev, Mikhail V. Gorshkov, Sergei A. Moshkovskii

**Affiliations:** Institute of Biomedical Chemistry, Moscow, Russia; Pirogov Russian National Research Medical University (RNRMU), Moscow, Russia; Karolinska Institutet, Stockholm, Sweden; Engelhardt Institute of Molecular Biology, Russian Academy of Sciences, Moscow, Russia; Institute of Energy Problems of Chemical Physics, Russian Academy of Sciences, Moscow, Russia; Moscow Institute of Physics and Technology (State University), Dolgoprudny, Moscow Region, Russia

**Keywords:** Proteogenomics, RNA editing, RNA-dependent adenosine deaminase, ADAR, shotgun proteomics, multiple reaction monitoring, SNARE complex, endophilin A

## Abstract

Adenosine-to-inosine RNA editing is one of the most common types of RNA editing, a posttranscriptional modification made by special enzymes. We present a proteomic study on this phenomenon for *Drosophila melanogaster*. Three proteome data sets were used in the study: two taken from public repository and the third one obtained here. A customized protein sequence database was generated using results of genome-wide adenosine-to-inosine RNA studies and applied for identifying the edited proteins. The total number of 68 edited peptides belonging to 59 proteins was identified in all data sets. Eight of them being shared between the whole insect, head and brain proteomes. Seven edited sites belonging to synaptic vesicle and membrane trafficking proteins were selected for validation by orthogonal analysis by Multiple Reaction Monitoring. Five editing events in *cpx, Syx1A, Cadps, CG4587* and *EndoA* were validated in fruit fly brain tissue at the proteome level using isotopically labeled standards. Ratios of unedited-to-edited proteoforms varied from 35:1 (*Syx1A*) to 1:2 (*EndoA*). Lys-137 to Glu editing of endophilin A may have functional consequences for its interaction to membrane. The work demonstrates the feasibility to identify the RNA editing event at the proteome level using shotgun proteomics and customized edited protein database.

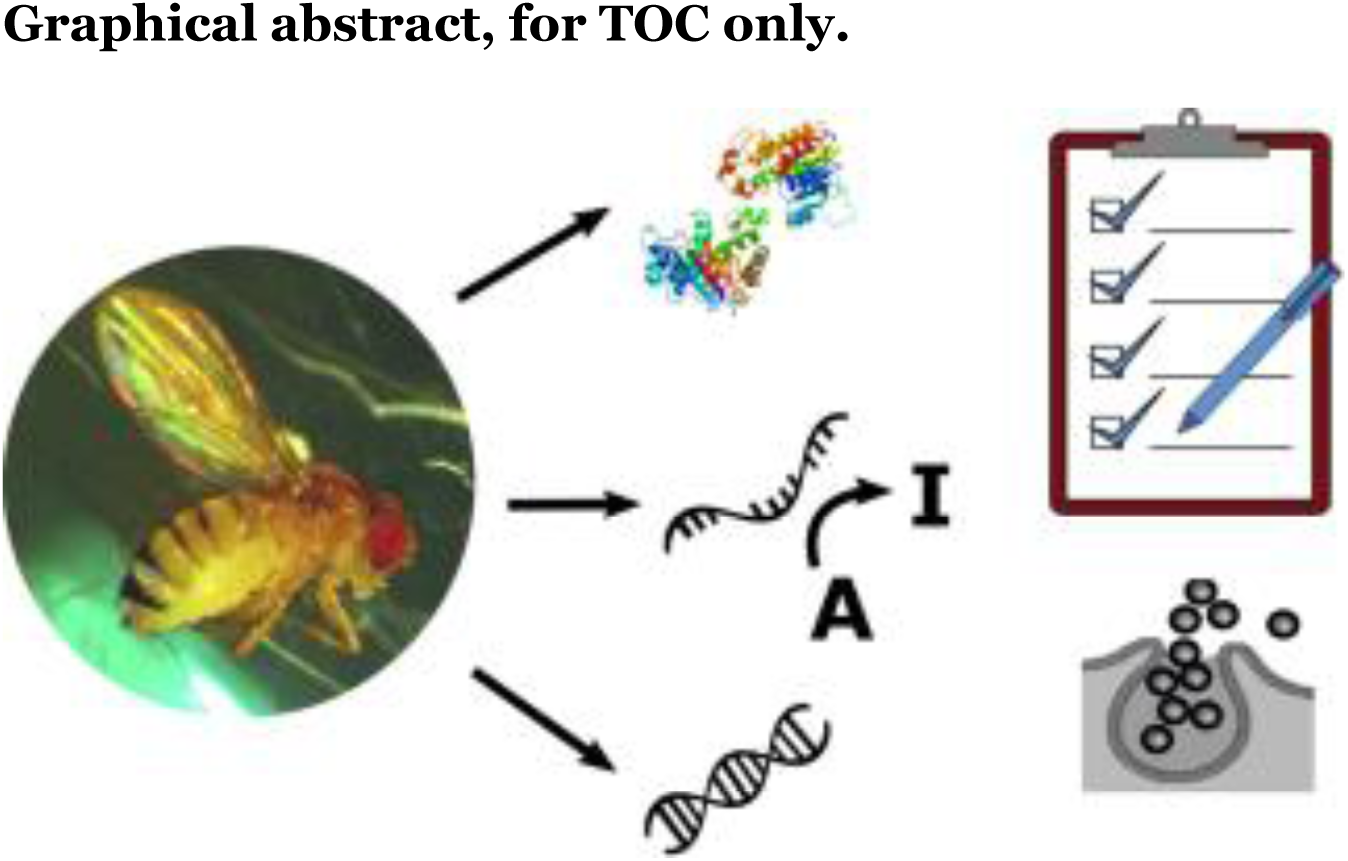

## Introduction

RNA editing is a type of posttranscriptional modification made by specific enzymes. Being first described to happen in mitochondrial RNAs of kinetoplastid protozoa ^1^, it was then observed for various organisms and different kinds of RNA ^2^. RNA editing includes nucleotide insertion or deletion, as well as deamination of cytosine and adenosine bases. Cytosine gets transformed into uridine by cytidine deaminase (CDA) ^3^, and adenosine is converted to inosine by adenosine deaminases acting on RNA (ADARs) ^4,5^. While the former is described mostly for plant cells ^6,7^, although it occurs also during apolipoprotein B synthesis ^3^, the latter is common for neural and glandular tissues of many invertebrate and vertebrate species ^8^.

Messenger RNA editing is the most interesting kind of RNA editing for proteomics as it may affect the primary structure of proteins. At the same time, specifically adenosine to inosine (A-to-I) editing provides the interest to neurobiology, because, reportedly, this type of modification is believed to have a function of rapid fine neuron tuning ^9^. Protein products of RNA editing can exist in organisms in both variants. The ratio of these proteoforms may possess a functional significance ^10,11^.

To date, the phenomenon of RNA editing was studied mostly at the transcriptome level that included a number of works on *Drosophila melanogaster* ^12^. Yet, a workflow for identification and characterization of the RNA editing products have not been fully developed. The first study on the RNA editing at the proteome level was focused on characterization of all types of proteoforms for the rat liver ^13^. However, liver is not reported as a tissue of functional A-to-I RNA editing ^14^, and twenty events of RNA editing identified for rat proteome were simply listed without further discussion ^13^.

After publishing the first version of the current work as a preprint, two more papers were represented that studied ADAR-mediated editing events at the level of proteome. First of them disclosed a lot of RNA editing sites in an octopus ^15^ and stated that this animal widely used RNA editing to tune its cellular functions in various environmental conditions. Flexibility of the cephalopod editome in comparison with mammalian species, at the same time, provided its genome conservation. In this work, which was mainly focused on RNA analysis, a survey of proteome also was done to identify hundreds editing sites. However, the authors did not use group-specific filtering of the results and those latter were not validated by orthogonal methods.

More recent paper has extracted proteins changed by RNA editing in human cancer tissues from proteogenomic big data of TCGA cancer project ^16^. In total, 13 editing sites were deduced and properly validated in proteomes from TCGA data. At least one of those sites, in *Copa* vesicle transport protein, was suspected to be involved in breast cancer progression^16^.

With the introduction of proteogenomic approach as use of customized nucleic acid databases for specific samples ^17^, the workflow for proteomic investigation of the products of RNA editing became pretty clear. First, a customized proteomic database is made, based on known editome of the organism under study ^18^. As the editome includes extra variants of the edited mRNA sequences, this database contains both unedited and edited peptide variants. Then, the shotgun proteomic spectra are searched against this database. Finally, information about the edited peptides is extracted from the search results and optionally validated to exclude false discoveries ^19^.

A-to-I RNA editing at the transcriptome level has been studied comprehensively for *D. melanogaster* by Hoopengardner et al. using comparative genomic approaches ^20^. In the other study by Rodriguez et al. authors used nascent RNA sequencing ^21^. Comparing wild type and the *adar* mutant flies they have shown the critical role the ADAR is playing in RNA editing. A method of cDNA to genomic DNA comparison was used to find RNA editing sites by Stapleton et al. ^22^.

More recently, a genome-wide analysis of A-to-I RNA editing sites was performed and an editome of *Drosophila* was thoroughly characterized ^2^3. The analysis revealed 3581 high-confidence editing sites in the whole body of a fruit fly. The authors used a single-molecule sequencing method with the introduction of the so-called ‘three-letter’ alignment procedure to avoid misreading of the A-to-G substitution sites for wild type and *adar*-deficient flies. This allowed increasing the accuracy of the database containing the A-to-I RNA editing sites and provided the most complete editome of *D. melanogaster*. This editome was further used for generating the customized protein database in this work. The summary of the results of previous efforts to study the *D. melanogaster* editome and the evolutional analysis of the function of A-to-I editing in seven *Drosophila* species was also provided last year by Yu et al. ^24^.

It also has been shown that A-to-I editing happens as a response to environmental changes such as temperature ^12^, which makes great sense in terms of the purpose of editing versus genomic recoding evolutionally. The authors have described 54 A-to-I editing sites, some of which are demonstrating significant differences in edited-to-unedited transcript ratios in the flies raised at 10, 20, and 30°C. The list of sites consists of various genes including *adar* itself.

From previous works with successful use of customized databases to identify protein-coding genome variants ^25^, we deduced that similar customized databases may be designed protein coding RNA editome. In this study, as a model animal we used a fruit fly with well characterized A-to-I RNA editome ^2^3. The aim of the study was identifying the RNA editing events in the proteome followed by validation of selected edited peptides by targeted mass spectrometry. To our knowledge, this is a first attempt to characterize RNA editing in *Drosophila* at the proteome level. Tandem mass spectrometry data were taken from recent shotgun proteomics studies ^26,27^ available at ProteomeXchange (http://www.proteomexchange.org/) ^28^. These results contain data for proteome of *Drosophila’s* whole bodies ^26^ and whole heads ^27^. The other data set for *Drosophila’s* brain proteome was obtained here using high-resolution mass spectrometry.

## Experimental section

### Experimental design

The shotgun proteomic analysis was performed using 1 sample consisted of 200 isolated fruit fly brains combined. The number of technical replicates in the shotgun experiment was 3. For the targeted proteomic experiment 2 samples had been prepared. The first one was used for preliminary MRM experiment and had been derived from flies of different age. Two RNA editing sites were validated in this experiment: *cpx* and *Syx1A*. During each MRM experiment 5 technical replicates have been done. The details of each experiment are provided below. The second MRM experiment was performed on the sample consisting of 80 brains of 72 hours old male flies. The sites validated in the second MRM experiment are *cpx, Syx1A*, CG4587, *Atx2, Cadps, RhoGAP100F*, and *EndoA*. A sample consisting of 100 fly heads was used for the genomic sequencing experiment and another 100 head sample was utilized for RNA study. The summarizing table of all the *Drosophila* samples used in this work is provided in Table S-1.

The data from 3 proteomes were used for the RNA editing sites search. One proteome was obtained experimentally here and the other two were taken from Xing et al. ^26^ and Aradska et al. ^27^. A thorough schematic explanation of the whole workflow performed is given in Fig. 1.

During the shotgun data analysis, the peptide identification was held at a 1% false discovery rate as described in details in the corresponding section.

**Figure 1.**
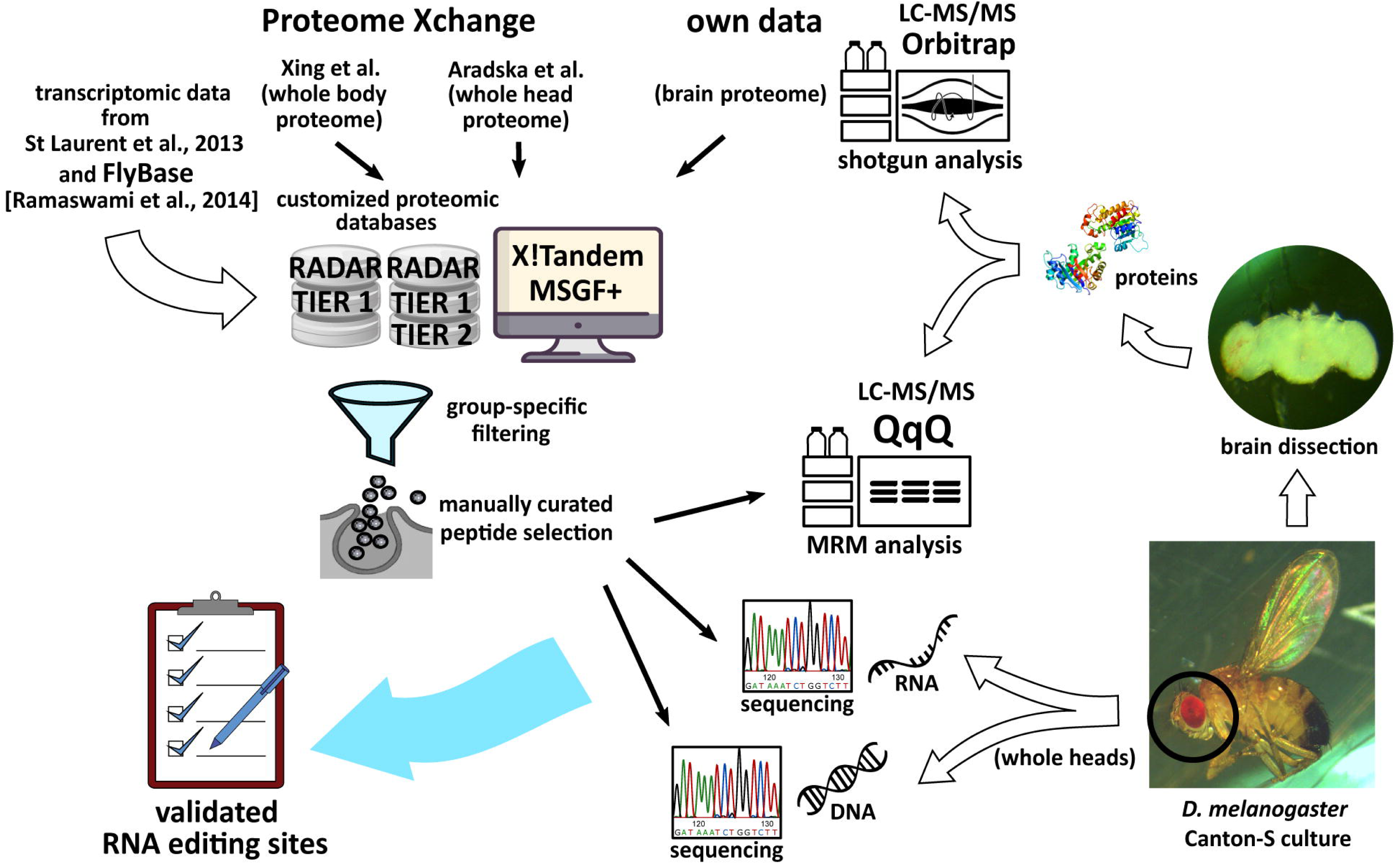
The workflow of this study. Fly brains were extracted and subjected to trypsin digestion following LC-MS/MS analysis. The heads were used for DNA and RNA extraction with the following genomic and transcriptomic sequencing. Two more datasets were taken from Proteome exchange from Xing et al. ^26^, and Aradska et al. ^27^ Two customized databases were generated and all three proteomic datasets were searched against them with two search engines. The transcriptomic data were taken from St Laurent et al. ^23^ and RADAR ^29^. Finally, a list of validated RNA editing sites was generated.

#### *Drosophila melanogaster* culture

Live samples of *Drosophila melanogaster* Canton S line were kindly provided by Dr. Natalia Romanova from Moscow State University, Department of Biology. The flies were kept on Formula 5-24 instant Drosophila medium (Carolina Biological Supply Company, USA) in 50 ml disposable plastic test tubes. The initial temperature for the fly culture was 25°C. The flies had been transferred to a new tube as they reached the adult stage. For general samples that were used for shotgun proteome analysis, as well as for the initial targeted analysis. Two hundred flies were selected from the culture regardless of their age. For the second MRM experiment 80 male flies were used, 72 hours old.

### Brain dissection

Frozen flies were kept on ice in a Petri dish during the whole procedure. Each fly was taken out, and the body was rapidly removed by a needle. The head was placed into 0.01 M PBS at pH 7.4 (recovered from tablets, Sigma-Aldrich, USA), and the head capsule was torn apart by two forceps under visual control through a stereo microscope (Nikon SMZ645, Japan) with 10×1 magnification. The extracted brains were collected into the same PBS, and then centrifuged at 6000 g for 15 minutes (Centrifuge 5415R, Eppendorf, Germany). The buffer solution was removed and the brain pellet was frozen at −80°C for the future sample preparation. The photo of the dissected brain was taken with DCM510 Microscope CMOS Camera (Scope Tek, China).

### Protein extraction, trypsin digestion, total protein and peptide concentration measurement

Brain pellet containing 200 brains was resuspended in 100 μL lysis solution containing 0.1 % (w/v) Protease MAX Surfactant (Promega. USA), 50 mM ammonium bicarbonate, and 10% (v/v) acetonitrile (ACN). The cell lysate was stirred for 60 min at 550 rpm at room temperature. The mixture was then subjected to sonication by Bandelin Sonopuls HD2070 ultrasonic homogenizer (Bandelin Electronic, Germany) at 30% amplitude using short pulses for 5 min. The supernatant was collected after centrifugation at 15.700 g for 10 min at 20°C (Centrifuge 5415R, Eppendorf, Germany). Total protein concentration was measured using bicinchoninic acid assay (BCA Kit, Sigma-Aldrich, USA).

Two μL of 500 mM dithiothreitol (DTT) in 50 mM triethylammonium bicarbonate (TEABC) buffer were added to the samples to the final DTT concentration of 10 mM followed by incubation for 20 min at 56°C. Thereafter, 2 μL of 500 mM iodoacetamide (IAM) in 50 mM TEABC were added to the sample to the final IAM concentration of 10 mM. The mixture was incubated in the darkness at room temperature for 30 min.

The total resultant protein content was digested with trypsin (Trypsin Gold, Promega, USA). The enzyme was added at the ratio of 1:40 (w/w) to the total protein content and the mixture was incubated overnight at 37°C. Enzymatic digestion was terminated by addition of acetic acid (5% w/v).

After the reaction was stopped, the sample was stirred (500 rpm) for 30 min at 45°C followed by centrifugation at 15,700 g for 10 min at 20°C. The supernatant was then added to the filter unit (10 kDa, Millipore, USA) and centrifuged at 13,400g for 20 min at 20°C. After that, 100 μL of 50% formic acid were added to the filter unit and the sample was centrifuged at 13,400 g for 20 min at 20°C. The final peptide concentration was measured using Peptide Assay (Thermo Fisher Scientific, USA) on a NanoDrop spectrophotometer (Thermo Fisher Scientific, USA). Sample was dried up using a vacuum concentrator (Eppendorf, Germany) at 45°C. Dried peptides were stored at −80°C until the LC-MS/MS analysis.

### Shotgun proteomic analysis

Chromatographic separation of peptides was achieved using homemade C18 column, 25 cm (Silica Tip 360μm OD, 75μm ID, New Objective, USA) connected to an UltimateTM 3000 RSLC nano chromatography system (Thermo Fisher Scientific. USA). Peptides were eluted at 300 nL/min flow rate for 240 min at a linear gradient from 2% to 26% ACN in 0.1% formic acid. Eluted peptides were ionized with electrospray ionization and analyzed on Orbitrap QExactive Plus mass spectrometer (Thermo Fisher Scientific, USA). The survey MS spectrum was acquired at the resolution of 60,000 in the range of m/z 200-2000.

MS/MS data for 20 most intense precursor ions, having charge state of 2 and higher, were obtained using higher-energy collisional dissociation (HCD) at a resolution of 15,000. Dynamic exclusion of up to 500 precursors for 60 seconds was used to avoid repeated analysis of the same peptides.

Proteomic data obtained in this work were deposited in the public repository ProteomeXchange (http://www.proteomexchange.org/) ^28^ under the accession number PXD009590.

### Customized database generation

Fly genomic coordinates of RNA editing sites mapped to exons were obtained from RADAR (1328 sites) ^29^ and genome-wide studies performed by St Laurent et al. ^23^. We used two lists of RNA editing sites from previous works ^23^. The first one, named TIER1 contained 645 exonic non-synonymous high-confident editing sites with high validation rate of >70%. The second list, named TIER2, contained 7986 less confident sites with expected validation rate of 9%. Genomic coordinates obtained from these sources were converted to the coordinates of the recent *Drosophila* genome assembly Dm6 using FlyBase (http://flybase.org/) ^30^. Changes in protein sequences induced by RNA editing were annotated for all three lists (RADAR, TIER1, and TIER2) using Variant Annotation Integrator (VAI) (http://genome.ucsc.edu/cgi-bin/hgVai). The input was prepared using Python script developed in-house (Script S-2). The VAI output was used to create VAI protein databases containing original and edited fly proteins using another Python script (Script S-3). These protein databases were used to generate the edited protein databases for MSGF+ and X!Tandem searches using another in-house developed Python script (Script S-4). The databases were named RADAR, TIER1, and TIER2 and can be found in Supporting information under the names Database S-5, S-6 and S-7, respectively.

### MSGF+ and X!Tandem search parameters

Peptide identification for all data sets was performed using MSGF+ version 2017.01.13 ^31^ as well as with X!Tandem version 2012.10.01.1 ^32^ against the three customized databases combined with the UniProt database for *D. melanogaster* downloaded in April 2017 and containing 42519 entries.

X!Tandem searches were performed using 10 ppm and 0.01 Da mass tolerances set for precursor and fragment ions, respectively. For the datasets obtained using a linear ion trap and taken from Xing at al.^26^, and Aradska et al.^27^, the fragment mass tolerance was 0.3 Da. Up to 1 missed cleavage was allowed. The same parameters were used for the MSGF+ search, except the instrument-specific parameters that were set according to the instruments specified for the particular data set. Carbamidomethylation of cysteine was used as fixed modification. Methionine oxidation and the protein N-terminal acetylation were used as variable modifications. The false discovery rate (FDR) was set to 1%. The group-specific FDR method, which provides separate FDR for the variant peptides as described elsewhere ^25^, was employed. A target-decoy approach was used to calculate FDR according to the following equation ^33^: FDR = (number of variant decoy PSMs+1)/number of variant target PSMs. All the edited peptides have been filtered group-specifically according to the 1% FDR. The filtration was performed using a Python script with a specific Pyteomics library constructed for he proteomic data analysis ^34^. Group-specific filtering was done separately for the peptides from RADAR, TIER1 and TIER2 databases.

### Open search

For open search, X!Tandem was used with the same settings except precursor mass tolerance which was set at 500 Da. The peptide-spectrum matches (PSMs) including decoys were grouped by mass shifts with the accuracy of 0.01 Da and filtered separately within each group. The open search was performed only on the data gained in our own experiment, since the other two data sets contain data with tandem spectra recorded in the linear ion trap with low resolution.

### Peptide standard synthesis

Peptides were synthesised by solid phase method using amino acid derivatives with 9-fluorenyl methyloxy carbonyl (Fmoc) protected α-amino groups (Novabiochem). The procedure was performed as described elsewhere^35^. Stable isotope-containing leucine (Fmoc-Leu-OH-^13^C6,^15^N, Cambridge Isotope Laboratories) was applied for labeling 11-plex peptides from *cpx* protein (NQMETQVNE***L^h^***K and NQIETQVNE***L^h^***K). A resin with attached stable isotope-labeled lysine (L-Lys (Boc) (^13^C_6_, 99%; ^15^N_2_, 99%) 2-Cl-Trt, Cambridge Isotope Laboratories) was used for synthesis of peptides of *Syx1A* (IEYHVEHAMDYVQTATQDT***K^h^*** and IEYHVEHAVDYVQTATQDT***K^h^***), *EndoA* (YSLDDNI***K****^h^* and YSLDDNIEQNFLEPLHHMQT***K^h^***), *Cadps* proteins (LMSVLESTLS***K^h^*** and LVSVLESTLS***K^h^***), and one of the peptides of *Atx2* (GVGPAPSANASADSSS***K^h^***). Resin with attached stable isotope-labeled arginine (L-Arg (Pbf) (^13^C_6_, 99%; ^15^N_4_, 99%) 2-Cl-Trt, Cambridge Isotope Laboratories) was used for synthesis of peptides of *CG4587* (LVTTVSTPVFD***R^h^*** and LVTTVSTPVFDG***R^h^***), *RhoGAP100F* (YLLQIWPQPQAQH***R^h^*** and YLLQIWPQPQAQHQ***R^h^***) and *Atx2* (GVGPAPSANASADSSS***R^h^***).

Further steps of synthesis were also preceded as described ^35^.

For synthesis quality control, a simple LC-MS analysis was held using a chromatographic Agilent ChemStation 1200 series connected to an Agilent 1100 series LC/MSD Trap XCT Ultra mass spectrometer (Agilent, USA). Since our peptides contained methionine residues, the quality control also included manual inspection of the MS and MS/MS spectra for possible presence of the peaks produced by oxidized compounds. No such peaks were found in our case.

Concentrations of synthesised peptides were determined using conventional amino acid analysis with their orthophtalic derivatives according to standard amino acid samples.

### Multiple Reaction Monitoring experiments

Each sample was analyzed using Dionex UltiMate 3000 RSLC nano System Series (Thermo Fisher Scientific, USA) connected to a triple quadrupole mass spectrometer TSQ Vantage (Thermo Fisher Scientific, USA) in five technical replicates. Generally, 1 μl of each sample containing 2 μg of total native peptides and 100 fmol of each standard peptide was loaded on a precolumn, Zorbax 300SB-C18 (5 μm, 5 × 0.3 mm) (Agilent Technologies, USA) and washed with 5% acetonitrile for 5 min at a flow rate of 10 μl/min before separation on the analytical column. Peptides were separated using RP-HPLC column, Zorbax 300SB-C18 (3.5 μm, 150mm × 75 μm) (Agilent Technologies, USA) using a linear gradient from 95% solvent A (0.1% formic acid) and 5 % solvent B (80% acetonitrile, 0.1% formic acid) to 60% solvent A and 40% solvent B over 25 minutes at a flow rate of 0.4 μl/minute.

MRM analysis was performed using Triple quadrupole TSQ Vantage (Thermo Scientific, USA) equipped with a nano-electrospray ion source. A set of transitions used for the analysis is shown in Table S-8. Capillary voltage was set at 2100 V, isolation window was set to 0.7 Da for the first and the third quadrupole, and the cycle time was 3 s. Fragmentation of precursor ions was performed at 1.0 mTorr using collision energies calculated by Skyline 3.1 software (MacCoss Lab Software, USA) (https://skyline.ms/project/home/software/Skyline/begin.view) (Table S-8). Quantitative analysis of MRM data was also performed using Skyline 3.1 software. Quantification data were obtained from the “total ratio” numbers calculated by Skyline represented a weighted mean of the transition ratios, where the weight was the area of the internal standard. Five transitions were used for each peptide including the isotopically labeled standard peptide. Isotopically labeled peptide counterparts were added at the concentration of 1 mg/ml. Each MRM experiment was repeated in 5 technical runs. The results were inspected using Skyline software to compare chromatographic profiles of endogenous peptide and stable-isotope labeled peptide. CV of transition intensity did not exceed 30% in technical runs.

All the MRM spectra can be downloaded from Passel (http://www.peptideatlas.org/passel/)^36^ under the accession number PASS01175.

### Genomic sequencing

DNA was extracted from 100 *Drosophila* heads (Table S-1, sample #5) using the standard phenol-chloroform method described elsewhere ^37^. The polymorphic sites of nine *D. melanogaster* genes (M244V in Syx1A, K398R in Atx2, Y390C in Atpalpha, R489G in CG4587, I125M in *cpx*, K137E in *EndoA*, Q1700R in *AlphaSpec*, M1234V in *Cadps* and Q1142R in *RhoGAP100F*) were genotyped using Sanger sequencing on Applied Biosystems 3500xL genetic analyzer and SeqScape® software (Thermo Fisher Scientific, USA). Initial PCRs were performed in a 25 μL volume containing 50 ng genomic DNA template, 10x PCR buffer, 0.5 U of HS Taq DNA Polymerase, 0.2 mM dNTPs (all from Evrogen, Russia), and 80 pmol of each primer. The PCR cycling conditions were the same for all SNPs and were as follows: 95°C for 5 minutes followed by 35 cycles of 94°C for 15 seconds, 59°C for 20 seconds, 72°C for 20 seconds and final elongation at 72°C for 6 minutes. Primers were designed using PerlPrimer free software (http://perlprimer.sourceforge.net/) and are shown in the Table S-9. The same primers were used for sequencing. PCR products were then cleaned up by incubation with the mix of 1 U of ExoI and 1 U of SAP (both enzymes from Thermo Fisher Scientific, USA) at 37° C for 30 minutes, followed by 80° C for 15 minutes. The sequencing reactions with following EDTA/ethanol purification were carried out using BigDye Terminator v3.1 Cycle Sequencing Kit (Thermo Fisher Scientific, USA) according to manufacturer’s instructions.

### RNA sequencing with inosine chemical erasing

In order to identify inosines on RNA strands, inosine chemical erasing (ICE) was used ^38^ with minor modifications. Briefly, total RNA from 100 fly heads was extracted using an RNeasy mini kit (Qiagen. Germany). Then, 10 μg of RNA was cyanoethylated (CE) by incubation in 38 μl solution (50% (v/v) ethanol, 1.1 M triethyl ammonium acetate, pH 8.6) with (CE+) or without (CE-) 1.6 M acrylonitrile at 70°C for 30 min. After cyanoethylation, RNA was purified using an RNeasy MinElute kit (Qiagen, Germany) and reverse transcribed using Low RNA Input Linear Amp Kit (Agilent Technologies, USA). Sites of nine *D. melanogaster* genes (*Syx1A, Atx2, Atpalpha, CG4587, cpx, EndoA, Alpha-Spec, Cadps, RhoGAP100F*) were analyzed by a Sanger sequencing described in the previous section. According to the known method ^38^, the editing site in CE- state can be detected as combined signal from A and G in the sequence chromatogram. For CE+ state, the signal can be detected as A. For those sites that are almost 100% edited, no amplification of the cDNA can be detected in the CE+ state ^38^.

## Results

### Search for RNA editing sites in deep *Drosophila* proteome

Tandem mass spectrometry data were taken from recent shotgun proteomics studies ^26,27^ available at ProteomeXchange. These results contain data for proteome of *Drosophila’s* whole bodies ^26^ and whole heads ^27^. The other dataset for *Drosophila’s* brain proteome was obtained here using high-resolution mass spectrometry. Note also that the proteome characterized by Aradska et al. ^27^ contains only membrane proteins as they were intentionally extracted during the sample preparation. In total, three data sets, representing proteomes of the whole body, the head, and the brain of *Drosophila* were available for the analysis in this study.

As noted above, the search for the RNA editing sites was performed using the proteogenomic approach ^39^. Following this approach, the standard fruit fly proteome FASTA database was extended with addition of the edited protein variants found from the transcriptomic data. Three FASTA files containing protein databases with edited sequences derived from transcriptome sequencing results ^23^ and FlyBase were generated as described in Method section and named TIER1, TIER2, and RADAR. These FASTA files were combined with the UniProt database for *D. melanogaster* (version form April of 2017, 42519 entries) containing the unedited peptides of fruit fly and used for the searches. After that, group-specific filtering was done separately for the peptides from RADAR, TIER1 and TIER2 databases. Figure 1 shows schematically the workflow used in this work. The edited peptide identifications are listed in Table 1. The genomic coordinates, unedited sequences and UniProt IDs of the peptides are listed in Table S-1.

**Table 1.**
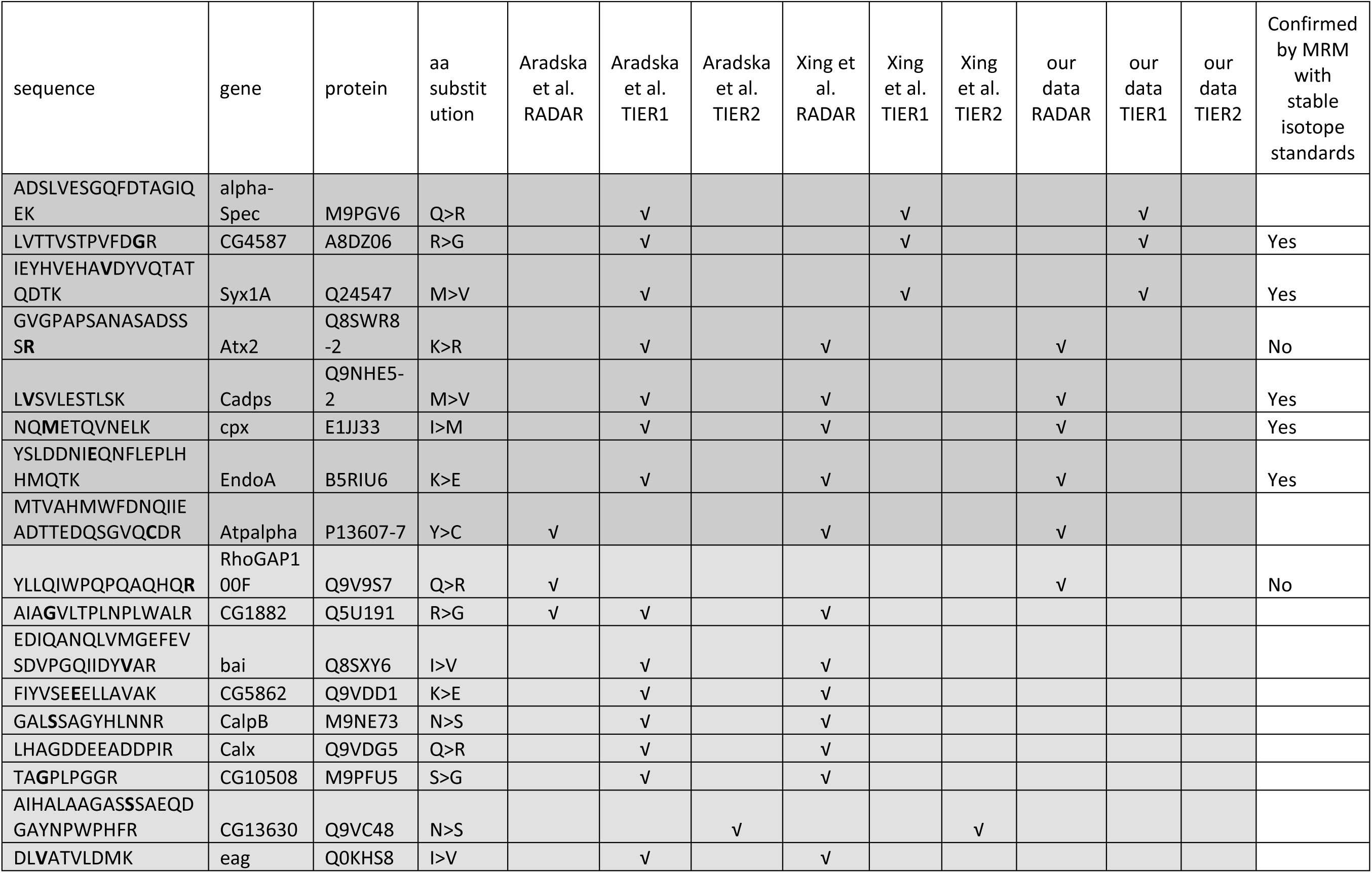
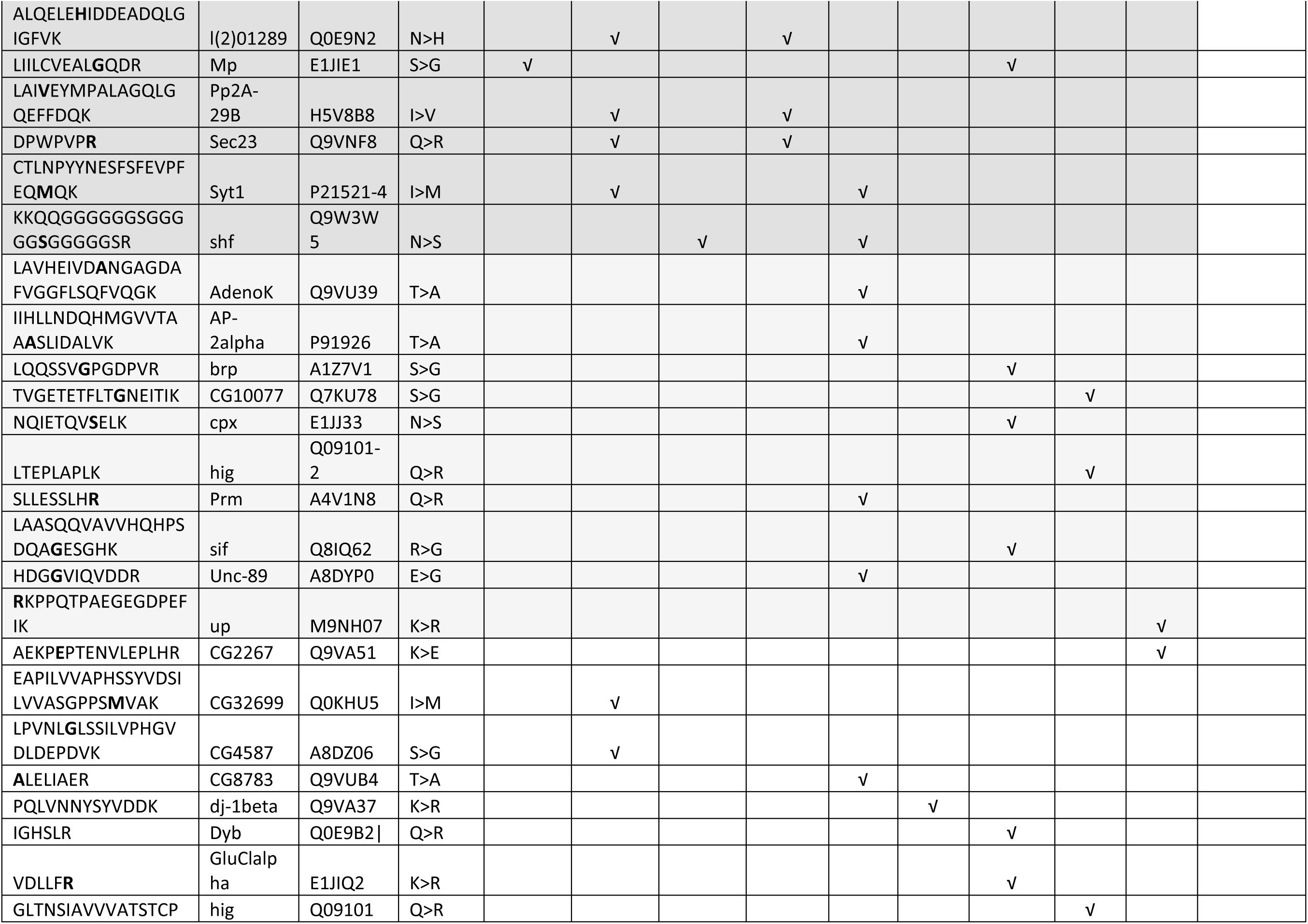
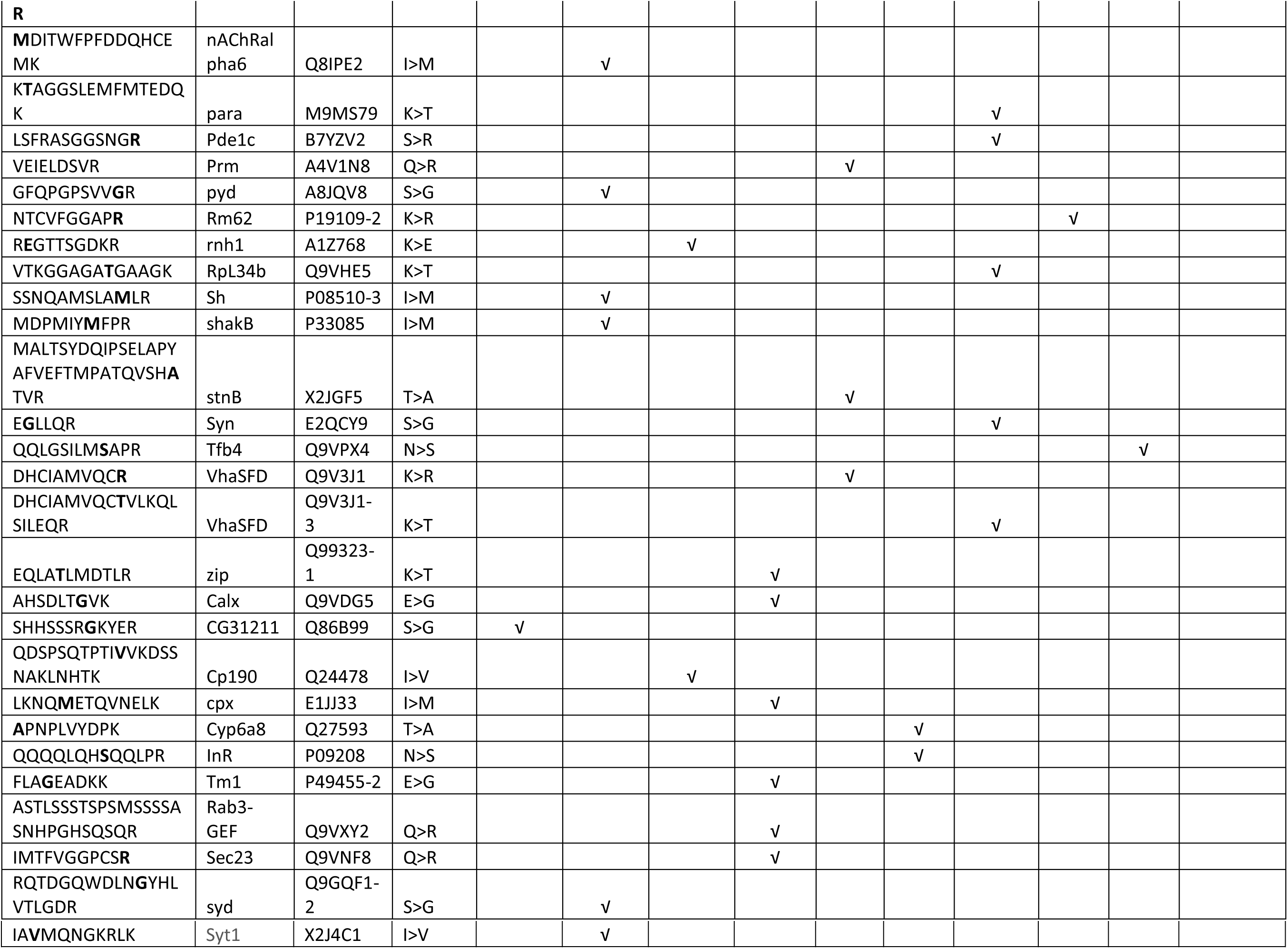
The edited peptides found in Aradska et al. ^27^, Xing et al.^26^, and in our data. The tiers of confidence are highlighted.

In all the data studied, 68 peptides corresponding to the RNA editing events were identified (Table 1). These peptides represent 59 proteins, since some of them, according to transcriptomic data, could contain several editing sites ^23^. However, because a mass-spectrometry provides partial protein sequence coverage only, some of the editing events for a protein are missing in the shotgun proteomics data.

Among edited sites listed in Table 1, only 8 are identified in all datasets used in the study. This group, as expected, includes proteins with brain localization, because in-home data is produced from isolated brains. Other 25 sites are found in at least two of datasets. Finally, remainder of edited sites (35) are found only in one study.

### Open search for additional filtering of false-positives

Open search was performed in order to assign some level of confidence to a particular amino acid substitution that is found to happen due to RNA editing, but may also be a false result introduced by an amino acid modification ^40^. The results of the open search are presented in Table S-10 (columns AA, AB, AC). Each amino acid substitution has its “open search rank” which was calculated based on percentage of open search hits falling into corresponding mass shift interval. If this mass shift would be overrepresented, most likely, it corresponded to the chemical modification rather than to the real amino acid substitution. Thus, the open search rank indirectly and inversely represents the likelihood of its mass shift to be caused by a modification. Based on the open search approach, some peptides containing N-to-D and Q-to-E substitutions have been removed from the results because such substitution cannot be told from deamidation. The open search has shown that some substitutions have higher likelihood to be mimicked by modifications than others.

### Functional annotation of edited proteins found in shotgun proteomes

To bare light on the purpose of RNA editing, all the peptides found to undergo editing shown in Table 1 were analyzed with the system of functional protein interactions (STRING, version 10.0 http://string-db.org/)^41^. Figure 2 shows the STRING analysis results. There are two groups of proteins with highly confident interactions. These groups were selected and named by manual curation based on Gene Ontology biological process analysis. In this part, we avoided use of software that automatically calculated enrichment by particular GO terms, since, in our case, it was difficult to determine the reference group of genes for this analysis.

**Figure 2.**
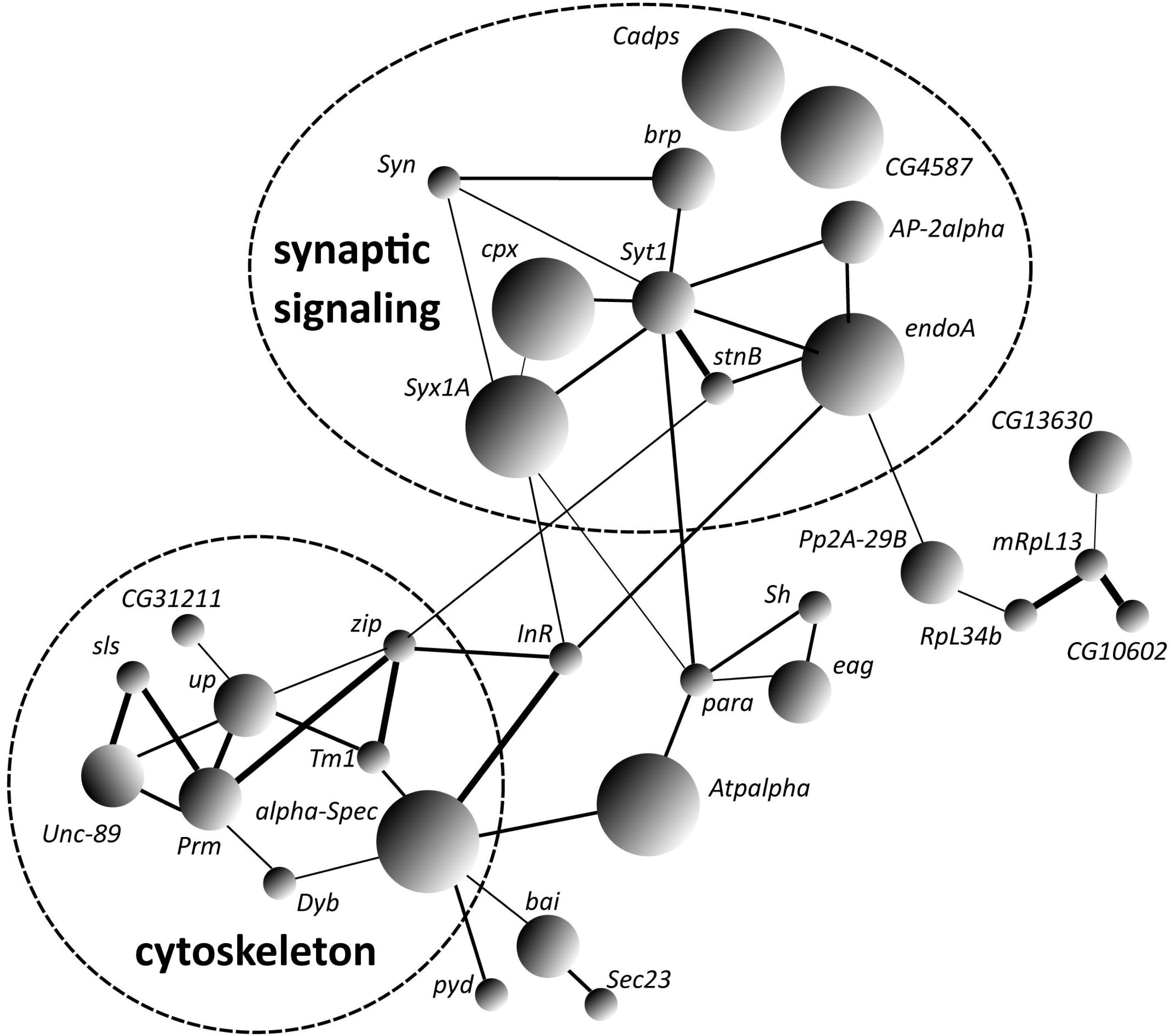
Interactions between proteins that undergo RNA editing. The size of the nods reflect the level of identification confidence in our analysis, the thickness of interaction lines shows the confidence of the interaction according to STRING.

First (“synaptic signaling” in the Fig. 2) group contains the following proteins: syntaxin (*Syx1A*), synaptotagmin (*Syt1*), complexin (cpx), synapsin (Syn), adaptin (*AP-2 alpha*), endophilin A (*endoA*), stoned protein B (*StnB*), Bruchpilot (brp), calcium-dependent secretion activator (*Cadps*), and calcium ion channel subunit encoded by *CG4587*. These proteins play a role in synaptic transmission. Particularly, syntaxin and synaptotagmin are components of a SNARE complex that provides fusion of synaptic vesicles with the presynaptic membrane, complexin being additional important binder of this complex ^42^. Stoned protein B also binds this complex and plays a role in synaptic vesicle endocytosis ^43^. Adaptins play a role in a process of synaptic vesicle recycling ^44^. Endophilin (*endoA*) acts in the process of vesicle endocytosis in neuromuscular junction ^45–48^. Calcium-dependent secretion activator (*Cadps*) is a Ca^2+^-dependent factor of vesicle endocytosis ^49,50^. The synaptic signaling group contains five proteins edited in all shotgun datasets, obviously representing most often edited protein products.

The second group (“cytoskeleton”) of proteins consists of non-muscular myosin (*zip*), alpha-Spectrin (*alpha-Spec*), titin (*sls*), and other gene products (Fig. 2). All proteins from this group are either components of cytoskeleton or interact with them and take a part in cell transport processes. This group includes only one edited site identified in all three datasets studied which belongs to alpha-spectrin protein.

### Genotyping of genomic DNA sites corresponding to found RNA editing events

Eight sites identified by shotgun proteome in all datasets used were further validated for A-to-I RNA editing at the level of nucleic acid. One site of *RhoGAP100F* gene found in two datasets was added to them due to close functional relation to other core edited proteins. Polymorphic sites of nine selected *D. melanogaster* genes (M244V in *Syx1A*, K398R in *Atx2*, Y390C in *Atpalpha*, R489G in *CG4587*, I125M in *cpx*, K137E in *EndoA*, Q1700R in *AlphaSpec*, M1234V in *Cadps* and Q1142R in *RhoGAP100F*) were genotyped. As a result, no traces of genetically encoded A-to-G substitutions were found which could probably mimic RNA editing events (Table 2). This finding confirms the assumption that the substitutions in nucleic acid, if they existed, happened post-transcriptionally.

**Table 2.** Validation of A-to-I editing sites at the level of genomic DNA, RNA and proteome. Sanger sequencing of amplified DNA was performed to confirm that the substitutions have happened post-transcriptionally. RNA editing events in the transcriptome level were confirmed using ICE method^38^. Edited protein sequences were validated using Miltiple Reaction Monitoring (MRM) using stable isotope labeled standards.

| Gene name | Putative genomic variant | Sequence of amplified DNA | Sequence of amplified cDNA |  | RNA editing confirmed by ICE-seq | Amino acid change confirmed by MRM |
| --- | --- | --- | --- | --- | --- | --- |
|  |  |  | CE- | CE+ |  |  |
| *Alpha-Spec* | A/G:Q1700R | T | A | A | No | N/A |
| *Atpalpha* | A/G:Y390C | T | A/G | A | **Yes** | N/A |
| *Atx2* | A/G:K398R | T | A | A | No | No |
| *Cadps* | A/G:M1234V | T | A/G | A | **Yes** | **Yes** |
| *CG4587* | A/G:R489G | T | A | A | No | **Yes** |
| *Cpx* | A/G:I125M | T | A/G | No product | **Yes** | **Yes** |
| *EndoA* | A/G:K137E | T | A | A | No | **Yes** |
| *RhoGAP100F* | A/G:Q1142R | T | A | A | No | No |
| *Syx1A* | A/G:M244V | T | A/G | A | **Yes** | **Yes** |

### Identification of inosine sites on RNA strands by inosine chemical erasing

Sites of nine genes selected as above were analyzed by inosine chemical erasing (ICE) method^38^. After comparison between cDNA sequencing from chemically erased (CE+) by acrylonitrile and intact (CE−) RNA, only in four of nine candidate sites, inosine-containing sites were confirmed by the chemical method including *Atpalfa* (chr3R:20965039:A/G:Y390C). *Cadps* (chr4:1265107:A/G:M>V). *Cpx* (chr3R:4297504:A/G:I125M) and *Syx1A* (chr3R:24103714:A/G:M244V) (Table 2).

### Targeted validation and quantitation of edited sites at the protein level by Multiple Reaction Monitoring (MRM) with stable isotope labeled standards

After shotgun analysis of a fruit fly editome on proteomic level a targeted analysis of the most interesting sites was held as a reasonable continuation of the study. We also intended to check the discrepancy between shotgun results and RNA study detected only five of nine sites of interest as inosine-containing.

For analysis, those sites present in all shotgun datasets were selected that were functionally related to synaptic function, more precisely, to pathways providing a fusion of the synaptic vesicle to the presynaptic cell membrane upon action potential ^51^. These presumably edited proteins included syntaxin 1A (*Syx1A*) and complexin (*Cpx*) known as the participant and the binder of the SNARE complex, respectively ^52^. Other proteins of interest included endophilin A (*EndoA*) providing membrane shaping in synapses^45^, *Cadps* and *CG4587* involved in calcium signalling in presynaptic zone ^49,53^ and a GTPase activator. *RhoGAP100F*, participating the organization of presynaptic dense zone ^54^. As mentioned above, this latter was added to the list for targeted analysis, in spite it was identified in two datasets only. Also, we added to the analysis a site of ataxin-2 (*Atx2*), well-studied protein with relevance to memory and other neural functions ^55^.

Currently, assays for targeted MS analyses are classified into three tiers, as described ^56^. According to this classification, we could define a present study as Tier 2. In our work, first, we studied modified peptides in non-human samples using MRM method. Also, it was not a clinical research, but used stable isotope labelled synthetic internal standards which did not undergo purification.

As it was mentioned above, not every candidate site checked was confirmed on the RNA level. However, we included four of such failed sites to the group for validation by MRM. As it was shown in Table 2, targeted test in two cases confirmed ICE-seq RNA data (*Atx2, RhoGAP100F*), where no traces of edited or intact peptides were detected in the brain hydrolysates. In contrast, a good evidence was provided by MRM for sites of *EndoA* and *CG4587* in both genome encoded and edited form, in spite no confirmation of these sites by RNA analysis. Stable isotope labelled standards of variant peptides for each site behaved similarly to naturally occurring products of the hydrolysed brain proteome in MRM tests (Fig. 3). This provided strong evidence of their chemical identity. Lack of detection of editing sites by inosine chemical erasing could be explained by low yield of desired products of the treatment for these certain sites. Otherwise, abundances of mRNA and protein products of the same gene are not always correlated, at least, if they are measured as a snapshot without temporal dependence ^57^. Hypothetically, RNA products of the genes of interest could be quickly destroyed during the procedures with animals. Finally, three sites were properly confirmed as edited in both transcriptome and proteome, which belonged to *Cadps, Cpx* and *Syt1* genes.

**Figure 3.**
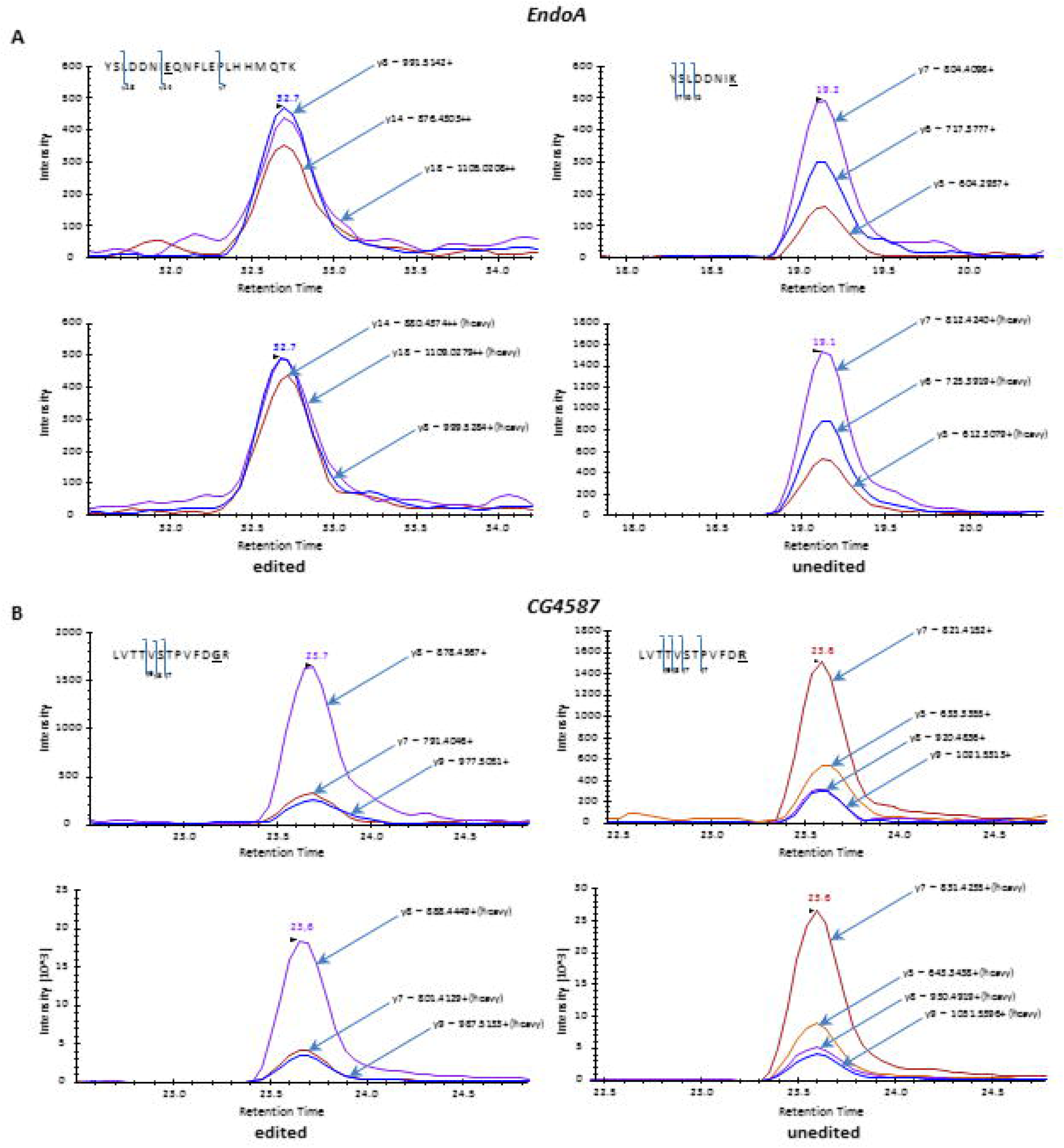
MRM spectra of edited and unedited variants of *CG4587* **(A)** and *EndoA* **(B)** peptides.

With labelled standards, we could measure absolute concentrations of targeted peptide in the brain samples. However, these levels may be influenced by many factors during multiple stages of sample preparation. According to our estimates, for five proteins of interest, these levels varied within one order of magnitude, from approximately 3.5 to 60 nmol/g of total protein in *Cadps* and *EndoA*, respectively (Fig. 4).

**Figure 4.**
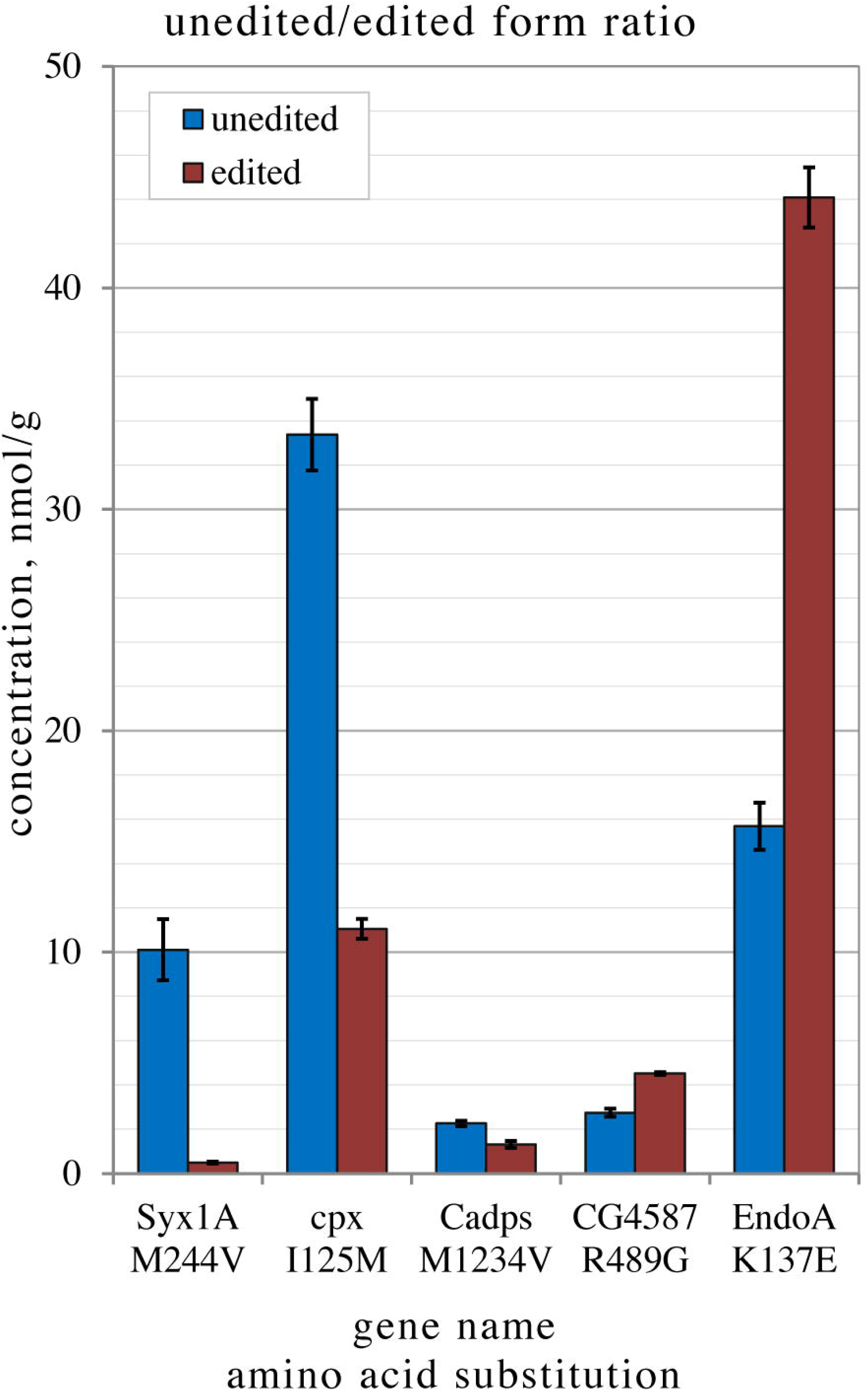
Concentrations of edited and unedited peptides of *Syx1A, cpx, Cadps, CG4587* and *EndoA* measured by MRM.

For aims of this work, it was more relevant to consider a ratio between genomic and edited forms of proteins which should be less dependent on sample preparation. This ratio was shown to be reproducible between two independent Canton-S fruit fly cultures (Figure S-11). At the same time, it varied between sites of interest. Thus, in syntaxin 1A, less than 5% of proteins occurred edited. Complexin, another member of SNARE complex was edited approximately by a quarter. *Cadps* product being edited by a third. Next, the *CG4587* calcium channel subunit and endophilin A sites were shown to be edited by 62% and 74%, respectively (Table 3). The latter protein attracted special attention, as it was simultaneously most abundant and most edited of products studied.

**Table 3.** Quantitation of selected edited sites and their genomic variants in the brain hydrolysate of the fruit fly by Multiple Reaction Monitoring (MRM) with stable isotope labelled standards.

| Gene name | Edited site | Tryptic peptide sequence for MRM study | | Result of ICE seq | Abundance, nmol/g total protein (av. ± st.d.) | CV, % | Ratio of edited site, % |
| --- | --- | --- | --- | --- | --- | --- | --- |
| *Syx1A* | M244V | Genomic | IEYHVEHAMDYVQTATQDTK | Yes | 10.1±1.4 | 13.7 | 4.6 |
| | | Edited | IEYHVEHAVDYVQTATQDTK | | 0.5±0.04 | 7.7 | |
| *cpx* | I125M | Genomic | NQIETQVNELK | Yes | 33.4±1.6 | 4.9 | 25.9 |
| | | Edited | NQMETQVNELK | | 11.1±0.5 | 4.1 | |
| *EndoA* | K137E | Genomic | YSLDDNIK | No | 15.6±1.1 | 6.9 | 73.8 |
| | | Edited | YSLDDNIEQNFLEPLHHMQTK | | 44.1±1.4 | 3.1 | |
| *CG4587* | R489G | Genomic | LVTTVSTPVFDR | No | 2.8±0.2 | 6.7 | 62.2 |
| | | Edited | LVTTVSTPVFDGR | | 4.5±0.1 | 1.5 | |
| *Cadps* | M1234V | Genomic | LMSVLESTLSK | Yes | 2.3±0.1 | 5.5 | 36.7 |
| | | Edited | LVSVLESTLSK | | 1.3±0.2 | 12.5 | |
| *Atx2* | K337R | Genomic | GVGPAPSANASADSSSK | No | Not detected |  |  |
|  |  | Edited | GVGPAPSANASADSSSR |  |  |  |  |
| *RhoGAP100F* | Q1142R | Genomic | YLLQIWPQPQAQHR | No | Not detected |  |  |
|  |  | Edited | YLLQIWPQPQAQHQR |  |  |  |  |

## Discussion

Shotgun proteomics after taking on board high-resolution mass-spectrometry gained success in identification and quantification of proteins as gene products, so called, master proteins. Today, LC-MS/MS analysis using high resolution mass spectrometry provides deep proteome covering about 50% of human genome from a single sample ^58,59^. This ratio is even higher in model organisms which may be characterized by more open genomes with higher numbers of expressed proteins ^60^. Correspondingly, a next aim of proteomics is to catalogue proteoforms of each protein. i.e. multiple protein species originated from one gene ^61,62^. Here we catalogued proteoforms of the fruit fly that originated from coding events of RNA editing by ADAR enzymes.

Expectedly, not every amino acid substitution predicted from RNA data could be detected. Partly, it is the limit of detection which is much higher in proteomics than PCR-based genomics. Moreover, many of edited RNA may not be translated at all. It is hypothesised that ADAR enzyme originally marked two-chained RNA in struggle versus viral genomes. Today, the immune functions are generally associated with ADAR1, at least, in mammalian cells^63^. Thus, many RNAs edited may lose their stability and eliminated without being translated.

Besides probably the original antiviral function, ADARs can recode mRNA and proteins, thereby changing their properties. A well-studied example is a Gln-to-Arg substitution in mammalian ion channel glutamate receptor *Gria2* which dramatically modulates their conductivity^64^. This protein recoding by ADAR better recognised in central neuron system and generally associated with ADAR2, at least, in mammals^63^. In our study, more likely, we encountered the functionally important recoding of proteins associated with synaptic function in the fruit fly.

Omics technologies are powerful instruments literally flooding researchers by qualitative and quantitative information. Thus, collecting transcriptome changes induced by its editing by ADARs, we checked their penetrance to the proteome level. However, we only can guess whether the changes identified are functional or represent a molecular noise as a side effect of the enzymatic activity. Although omics results are descriptive and cannot by themselves provide biological findings, they represent a good tool to hypothesize. Further we try to discuss the hypotheses that may result from validated proteomic data of ADAR-induced editome.

As mentioned above, syntaxin 1a is a component of the well-studied SNARE complex which provides fusion of presynaptic membrane and synaptic vesicles in most organism with neural system. SNARE is characterized structurally many times, especially, for model mammals^65^. In the complex, α-spirals of four proteins including syntaxin form a bunch. The editing site found in syntaxin in this work, M244V, is situated, however, outside this spiral and is not mapped to any known functional site. The importance of this specific site is dubious due to the lesser extent of its editing which was not above 5%.

Complexin is also a protein functionally related to SNARE complex. By binding to the bundle of α-spirals of the complex by its own spiral it prevents spontaneous fusion of vesicles with the membrane in absence of calcium signaling^66^. The I125M site, however, is located in the C-terminus of the molecule which is not conservative between rodents and insects. In rat, this part of complexin molecule provides membrane binding^66^. Hypothetically, the substitution caused by editing in complexin of the fruit fly may modulate affinity of protein-membrane interaction.

The Calcium-Dependent Secretion Activator (CADPS, or CAPS) expressed by *Cadps* gene acts from the other side of SNARE complex being connected with synaptic vesicles and activating them in presence of calcium ions^49^. CAPS was originally proposed to bind the SNARE complex after priming synaptic vesicles, but later its mechanism of action was shown to be SNARE-independent^67^. We have shown that more than a third of this protein was subjected to M1234V editing. However, this substitution could be not interpreted from the position of spatial structure or domain function. Notably, human and mouse orthologs of *Cadps* are known to be extensively edited by ADAR enzyme. According to RADAR database, each of these genes is edited in tens sites, at least two of them coding amino acid substitutions^29^. These latter are although not overlap the site edited in the fruit fly.

The product of the gene *CG4587* is a protein engaged in Ca^2^+-dependent nociception response ^53^. Little is known about its spatial structure, such that it is difficult to propose a role of the R489G substitution. There is some parallel of this type of substitution with the mammalian editome where glutamate ion channel subunits (*Gria2, Gria4*) also bear a similar amino acid change ^64^. Moreover, as in the case of CAPS protein, a human ortholog of *CG4587* called a voltage-dependent calcium channel subunit alpha-2/delta-4 (*CACNA2D4*) also is edited, although in the intronic region ^68^. Notably, based on our data, yet undiscovered importance of this gene product could be deduced. *CG4587* protein shares about 30% identity with its important paralog, Straitjacket (*stj*), which is another Ca^2^+-dependent channel subunit, also involved in nociception ^69^. Straitjacket is not in our list of the proteins corresponding to the RNA editing event, but it was found in the fruit fly proteome and identified by search engines in our data. Theoretically, *stj* may correspond to the editing event, because its transcript was prone to editing by ADAR and the protein was in our customized database (Supplemental files 5 and 6). The fact that *stj* did not listed among the identified edited proteins could be explained by its underrepresentation in the spectra compared to *CG4587*. The head and brain proteomes are of the most interest in terms of the expression of the Ca^2^+-dependent channels. In the brain proteome, the relative intensity of *stj* calculated by MaxQuant search engine is 34.7 times lower compared with *CG4587*. Label-free quantification (LFQ) obtained for X!tandem search results has shown the following results for *CG4587/stj* LFQ ratios: 12.4, 4.7 and 6.6 for SIN ^70^, NSAF ^71^, and emPAI ^72^ LFQ algorithms, respectively. For the whole head proteome, the *CG4587/stj* ratios were 5.4, 4.4, and 8.5 for these LFQ algorithms, respectively. Therefore, even though straitjacket is an important and well-characterized protein of nervous tissue, its paralog, *CG4587* gene product had significantly higher abundance in the proteomes of isolated brain, as well as the whole head. Along with the Straitjacket protein, this subunit deserves further functional studies.

Unlike of all other sites of interest which lacked structural information, we were luckier with endophilin A editing site, where lysine-137 was substituted to glutamate with a high yield of 74% (Table 3). This residue is located inside the α2-spiral of conserved and well-studied BAR domain of this protein ^73^. Domains of this type form dimers and are contained many proteins associated with intercellular membrane dynamics ^74^. It was shown that BAR domain in endophilin A provided the membrane curvature binding hydrophilic surface of lipid bilayer by concaved surface of its dimer ^75^. A residue of interest is located close to this surface or on it, as it can be seen from the spatial structure of human protein, which in this part is almost identical to that of *Drosophila* (Figure 5). The affinity between membrane and endophilin A BAR domain is provided *inter alia* by electrostatic interactions between negatively charged phospholipid heads and lysine residues on the concave part. Thus, one can hypothesise that a recharging Lys-to-Glu substitution in endophilin A may dramatically influence the binding affinity of the protein with membrane, whereby regulating membrane dynamics in neural cells. This finding can be verified by experiments with recombinant proteins and model membranes. The impact of endophilin A editing on the insect organism may be verified by generation of corresponding mutant strains.

**Figure 5.**
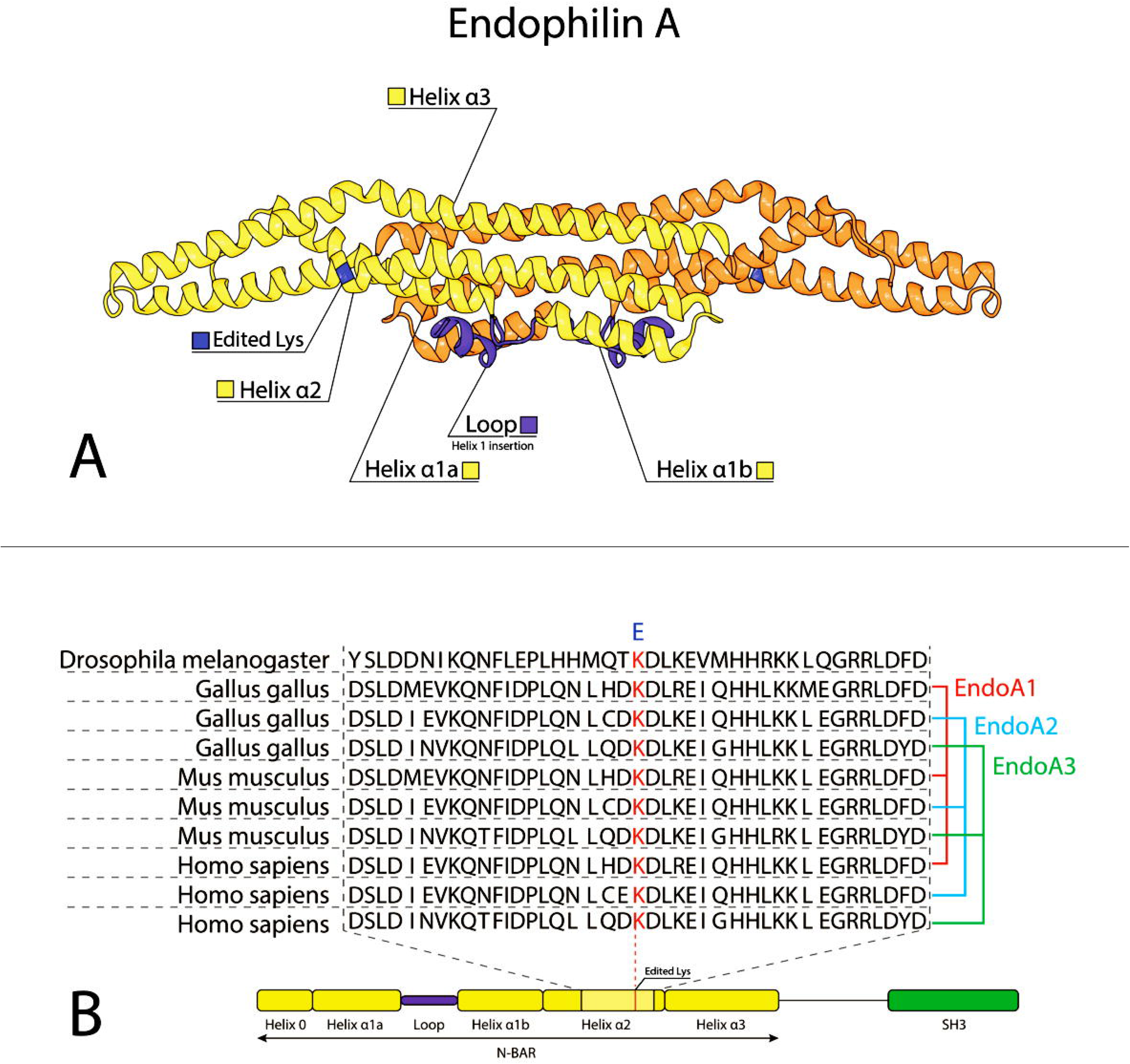
A lysine edited to glutamic acid in fruit fly endophilin A in the spatial structure of BAR domain. **A**. Homologous position of conservative lysine residue corresponding to Lys-137 of fruit fly *EndoA* shown in the dimer of BAR domains of the human ortholog (PDB accession 1X03) [Masuda et al.] ^74^ Structural elements of BAR domain are designated as described [Weissenhorn] ^72^. **B**. Local alignments between fruit fly *EndoA* product and endophilin A isoforms of chicken, mouse and human illustrate a high level of conservation in α2 helix between animals.

Notably, endophilin A transcripts in mammals are shown to be edited by ADAR enzymes, but in their intronic parts. In the contrary, for the fruit fly we observed massive recoding of the protein sequence, which, obviously, should have a functional impact. Recently, it was suggested that A-to-I editing in *Protostomia* could tune functions of proteins instead of genomic mutations^15^. What if the *endoA* editing in the fruit fly was functionally compensated in vertebrates by evolutional creation of three gene paralogs for endophilin A?

In this work, we identified consequences of adenosine-to-inosine RNA editing in shotgun proteomes of *Drosophila melanogaster*, validated and quantified selected edited sites by targeted proteomics of the brain tissue. Some of proteins with known neural function, for example, endophilin A, were shown to be substantially edited in the insect brain. Thus, our omics experiments provided some sound and testable hypotheses, which could be further verified by orthogonal biochemical methods.

## Acknowledgements

The work was funded by the Russian Scientific Foundation, grant # 17-15-01229.

Authors thank Dr. Natalia Romanova from the Department of Genetics of Biological Faculty of Moscow State University for providing the *Drosophila melanogaster* Canton S stock and Dr. Leonid Kurbatov from the Institute of Biomedical Chemistry for technical assistance.

